# Genetic variants drive altered epigenetic regulation of endotoxin tolerance in BTBR macrophages

**DOI:** 10.1101/2020.02.08.940296

**Authors:** Annie Vogel Ciernia, Verena M. Link, Milo Careaga, Janine LaSalle, Paul Ashwood

**Affiliations:** Department Biochemistry and Molecular Biology, Djavad Mowafaghian Centre for Brain Health, University of British Columbia, Vancouver, Canada V6T2A1; Metaorganism Immunity Section, Laboratory of Immune System Biology, National Institute of Allergy and Infectious Diseases, National Institutes of Health, Bethesda, MD USA 20892; Department of Medical Microbiology and Immunology, MIND Institute, Genome Center, University of California, Davis, CA USA 95616

## Abstract

The BTBR T^+^Itpr3^tf^/J (BTBR) mouse has been used as a complex genetic model of Autism Spectrum Disorders (ASD). While the specific mechanisms underlying BTBR behavioral phenotypes are poorly understood, prior studies have implicated profound differences in innate immune system control of pro-inflammatory cytokines. Innate immune activation and elevated pro-inflammatory cytokines are also detected in blood of children with ASD. In this study, we examined how underlying BTBR genetic variants correspond to strain-specific changes in chromatin accessibility, resulting in a pro-inflammatory response specifically in BTBR bone marrow derived macrophages (BMDM). In response to repeated lipopolysaccharide (LPS) treatments, C57BL/6J (C57) BMDM exhibited intact endotoxin tolerance. In contrast, BTBR BMDM exhibited hyper-responsive expression of genes that were normally tolerized in C57. This failure in formation of endotoxin tolerance in BTBR was mirrored at the level of chromatin accessibility. Using ATAC-seq, we specifically identified promoter and enhancer regions with strain-specific differential chromatin accessibility both at baseline and in response to LPS. Regions with strain-specific differences in chromatin accessibility were significantly enriched for BTBR genetic variants, such that an average of 22% of the differential chromatin regions had at least one variant. Together, these results demonstrate that BTBR genetic variants contribute to altered chromatin responsiveness to endotoxin challenge and a failure in formation of tolerance, resulting in a hyper-responsive innate immunity in BTBR. These findings provide evidence for an interaction between complex genetic variants and differential epigenetic regulation of innate immune responses. Our findings also provide novel mechanistic insight into the complex genetic architecture and immune abnormalities observed in ASD.

## Introduction

The ability of peripheral innate immune cells to control pro-inflammatory cytokines during periods of repeated or prolonged exposure to immune activating stimuli is critical for preventing tissue damage(Liu, 2019). Cells that compose the innate immune system such as monocytes, macrophages, natural killer cells and microglia were traditionally thought to respond rapidly and nonspecifically to immune stimuli and lack immune memory(Netea et al., 2011, 2016; Wendeln et al., 2018). However, recent evidence has demonstrated a form of innate immune memory in which immune activation can alter a secondary response to either the same or different pathogen exposure. In macrophages, repeated exposure to lipopolysaccharide (LPS) induces a refractory period of immune activation termed endotoxin tolerance. This form of innate immune memory results from epigenetic suppression of genes associated with inflammation (tolerance) and continued expression of genes coding for anti-microbial molecules (non-tolerance)(Foster et al., 2007; Novakovic et al., 2016; Saeed et al., 2014). Mechanistic studies have demonstrated that innate immune memory is critically dependent on epigenetic reprograming that produces transcriptional changes in immune signaling, metabolism and ultimately enhances the immune cell’s capacity to appropriately respond to stimulation(Novakovic et al., 2016; Saeed et al., 2014). How these epigenetic regulatory mechanisms become dysfunctional in cases of chronic inflammation is of great interest considering the large number of disease states characterized by increased inflammation.

Numerous studies have shown increased innate immune activation in individuals with autism spectrum disorders (ASD)(Ashwood et al., 2011; Enstrom et al., 2010; Goines and Ashwood, 2013; Krakowiak et al., 2017; Zerbo et al., 2014). These include increased activated monocyte populations circulating in the periphery or increased cytokines related to monocyte/macrophage activation(Hughes et al., 2018). For example, increased IL-1β was found in newborn blood spots from infants that later went on to develop ASD compared to typically developing children(Krakowiak et al., 2017). Monocytes isolated from children with ASD compared to typically developing children show elevated baseline cytokine levels, as well as a hyper-response in cytokine levels that were associated increased behavioral impairments in the children(Enstrom et al., 2010). Several additional lines of evidence point to increased microglia activation in ASD brain, including activated microglial morphology(Morgan et al., 2010, 2012, 2014), enhanced pro-inflammatory cytokines(Smith et al., 2012; Tarkowski et al., 2003), increased immune gene expression(Gandal et al., 2018a, 2018b; Parikshak et al., 2016), and real-time PET scan analyses(Suzuki et al., 2013).

In animal models relevant to ASD, elevated innate immune responses have also been shown(Onore et al., 2013; Schwartzer et al., 2013, 2015). The BTBR T^+^Itpr3^tf^/J (BTBR) mouse strain shows a natural lack of sociability, increased repetitive behaviors, and impaired cognition(McFarlane et al., 2008; Moy et al., 2007; Roullet and Crawley, 2011; Scattoni et al., 2011). Similar to monocytes isolated from children with ASD, macrophages from the BTBR mouse strain have a skewed pro-inflammatory phenotype and show a larger cytokine response to LPS challenge than macrophages from C57BL/6J (C57) mice(Onore et al., 2013). The immune alterations in BTBR have been correlated with the presentation of ASD relevant behaviors(Onore et al., 2013), suggesting a potential link between immune and behavioral dysfunction. Consistent with this hypothesis, replacement of the immune system with wildtype bone marrow rescued the social impairments but not the repetitive behaviors in BTBR(Schwartzer et al., 2017). BTBR mice were originally derived in the 1950s from an inbred strain carrying the a^t^ (nonagouti; black and tan) and wildtype T (brachyury) mutations and then subsequently crossed with mice with the tufted (Itpr3^tf^) allele (http://jaxmice.jax.org/strain/002282.html). The BTBR genome is divergent from that of C57, including over 6 million SNPs and 1.6 million insertions and deletions (InDels)(Jones-Davis et al., 2013; Pstkov et al., 2005), similar to differences between any two human individuals(Auton et al., 2015). In combination with the observed immune abnormalities, the BTBR mouse strain is an interesting model for ASD that captures aspects of complex genetic, environmental, and immune components of the disorder.

To uncover mechanisms that may explain BTBR genetic differences in regulation of cytokine responses, we examined gene expression and chromatin dynamics in response to immune challenge and tolerance using BMDM cultures from BTBR compared to C57 mice. We specifically assessed endotoxin responses in BMDM cultures(Foster et al., 2007) from BTBR and C57 strains to test the hypothesis that the chronic hyper-inflammatory state observed in BTBR mice may be a result of altered chromatin dynamics and immune gene regulation. Using a combination of gene expression, epigenetic, genetic, and bioinformatic approaches, we identified key regions of differential chromatin regulation between the strains that related to both altered gene expression patterns and underlying strain genetic variants. Together our findings indicate that genetic variation between strains significantly contributes to altered chromatin accessibility and LPS-induced gene expression in BMDM cultures, culminating in higher baseline and LPS responsivity in the BTBR strain.

## Results

### BTBR BMDM hyper-respond to LPS immune challenge

To test the hypothesis that BTBR strain differences in inflammation can be observed at a level of gene responsivity to LPS challenge and tolerance, we compared transcript levels of cultured bone marrow derived macrophages (BMDM) from juvenile postnatal day 28 (P28) male BTBR and C57BL6/J (C57) mice. After seven days in culture the differentiated macrophages were treated with either media, one or two doses of LPS (100 ng/ml) (Figure 1). This protocol has been utilized previously to examine endotoxin tolerance in cultured BMDM(Foster et al., 2007). We confirmed the expression patterns of gene targets involved in forming endotoxin tolerance using a custom Fluidigm Biomark HD Delta gene assay containing gene targets previously identified(Foster et al., 2007) as either tolerized (reduced expression upon the second LPS exposure) or non-tolerized (fail to suppress expression upon the second LPS exposure) (Figure 2 and Table S1).

**Figure 1.**
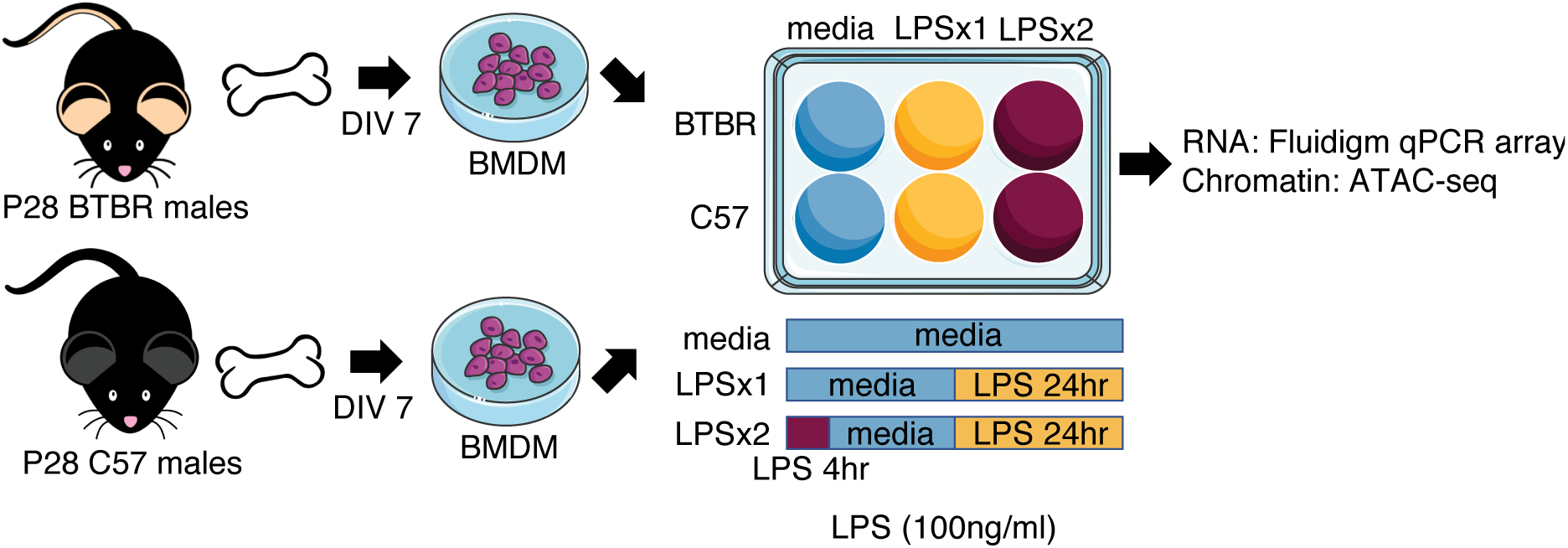
Experimental Design. Bone marrow was extracted from postnatal day 28 (P28) male BTBR or C57 mice and differentiated in culture for seven days (DIV 7) to form bone marrow derived macrophages (BMDM). BMDM from each strain were re-seeded into six well plates and treated with either media, one dose of LPS for 24hrs (LPSx1) or given a 4 hour LPS pretreatment followed by a 24hour LPS treatment (LPSx2). All LPS treatments were at 100ng/ml of culture media. Following treatment, cells were collected for RNA expression analysis (Fluidigm array) or chromatin accessibility (ATAC-sequencing).

**Figure 2.**
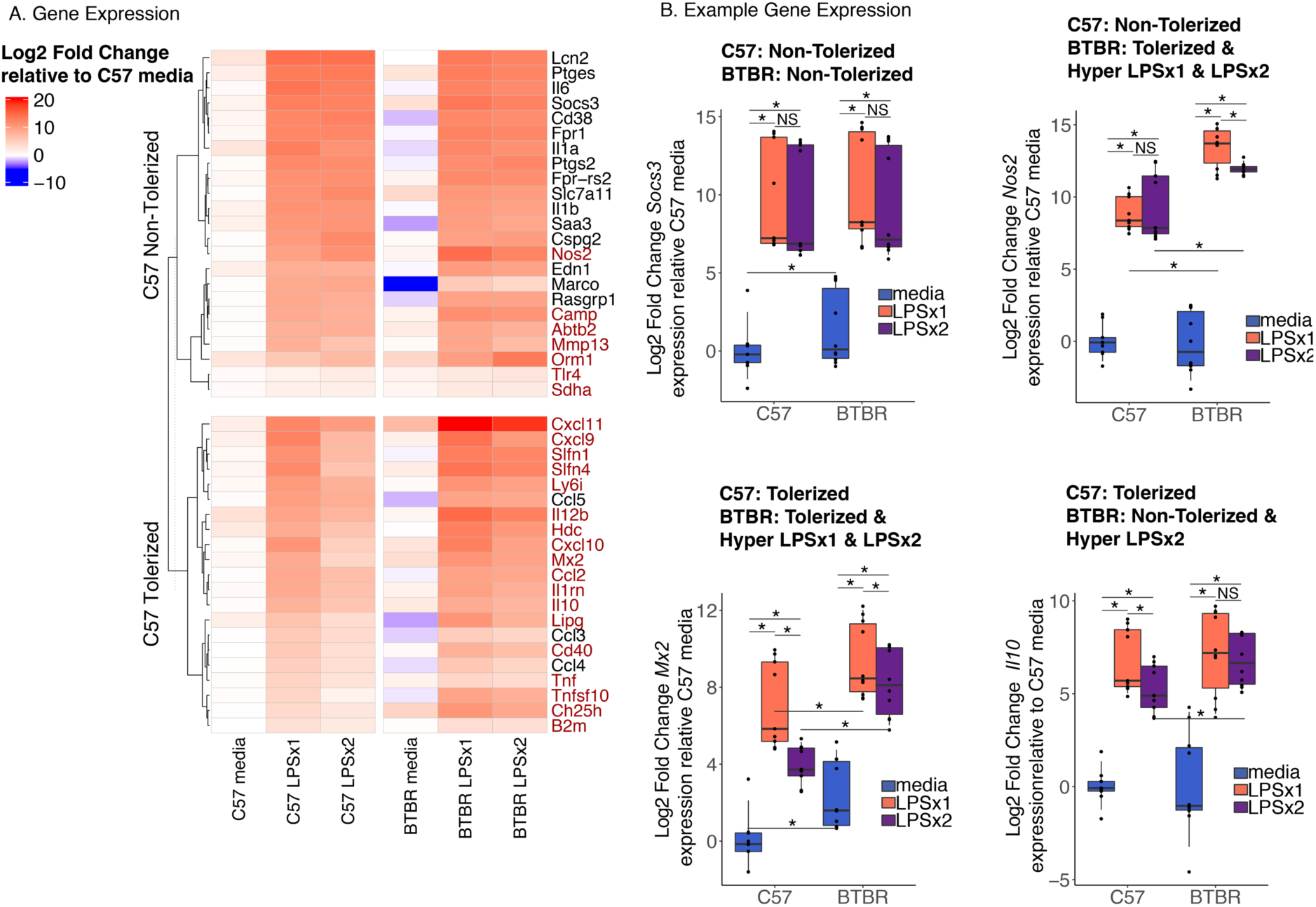
BTBR BMDM Hyper-Respond to LPS Stimulation. **A.** Log2 Fold Change deltadeltaCt relative to average C57 media values for genes on the Fluidigm qPCR expression array that show either non-tolerized or tolerized patterns of expression in C57 BMDM. Gene expression patterns were classified based on Tukey HSD corrected posthoc comparisons following a linear regression analysis (see methods) (Table S1). Genes highlighted in red show a significantly larger response in BTBR compared to C57 for LPSx1 or LPSx2, respectively (Tukey posthoc corrected comparisons p < 0.05). n=4-6 per condition. **B.** Boxplots of individual example genes for log2 fold change gene expression. Boxes are defined by the 25th and 75th quartiles for each condition with median values shown as the horizontal line. Whiskers are defined by the 5^th^ and 95^th^ percentile values. Individual values are shown as overlaid dots. * Tukey HSD corrected posthoc comparison *p*-value < 0.05. Categorizations given above each plot. See Table S1 for statistics.

In the C57 BMDMs, 21 of the 47 genes on the array showed an expression pattern consistent with tolerization, defined as an initial increase from baseline (media) with a single dose of LPS (LPSx1) and an attenuated response when the cultures had been pretreated with LPS (LPSx2) (Figure 2 and Table S1). Of the 47 genes on the array, 23 showed a non-tolerized pattern in which the pre-treatment with LPS failed to produce an attenuation of the second response (LPSx1 ∼ LPSx2). In addition, one gene (*Cd14*) showed a sensitized response with the pre-treatment of LPS producing a larger response after LPSx2 compared to LPSx1 (Table S1). In contrast, for the BTBR BMDMs, only 14/47 genes showed a tolerized pattern of expression, 32/47 showed a non-tolerized pattern, and one showed sensitization. Many of the examined genes (28/47) showed significant but small magnitude strain-specific differences in media condition (adjusted *p*-value < 0.05) (Figure 2, Table S1), suggesting that strain alone influences baseline regulation of some immune genes similar to what has been observed previously for other strains(Link et al., 2018a).

For genes showing a non-tolerized pattern of expression in C57, all but three showed a similar non-tolerized pattern in BTBR (see Figure 2B *Soc3* expression for an example). For genes that showed a tolerized expression in C57, 11 showed a similar tolerized expression in BTBR, in contrast,10 showed a non-tolerized pattern specifically in BTBR. Consistent with previous observations at the protein level in the BTBR mice, many of the examined genes showed a hyper-response to LPS treatment (labeled red in Figure 2A heatmap). For C57 non-tolerized genes, 7 out of 23 genes showed a larger response to either a single or multiple LPS treatments in the BTBR compared to C57 BMDM (see Figure 2B *Nos2* expression for an example). Even more striking was that for the 21 C57 tolerized genes, 85% (18/21) showed a significant (*p*-value < 0.05) hyper-response in BTBR with either one or both LPS treatments. For example, both *Mx2* and *Il10* showed significantly larger LPS inductions in BTBR compared to C57 (Figure 2B). Together, these results suggest that BTBR shows increased responsivity to LPS and that is particularly true of genes that normally become tolerized to LPS in C57.

### Chromatin accessibility across strains in response to LPS

To test the hypothesis that strain-specific differences in chromatin accessibility may dictate the transcriptional response profile observed in the BTBR BMDM, we performed Assay for Transposase-Accessible Chromatin using sequencing (ATAC-seq)(Buenrostro et al., 2013) on C57 and BTBR BMDMs stimulated with media, LPSx1 or LPSx2 treatments. There were no differences in alignment, any of the quality control (QC) metrics or in the number of filtered or unfiltered reads between any of the sample conditions (Table S2 and Figure S1). Two samples failed final QC assessment and were excluded from subsequent analysis (Figure S2 and S3). Differential regions of accessibility for ATAC-seq were identified within nucleosome free regions (NFR, reads <147bp) for each biological replicate and consensus peaks that were found in at least 3 out of 4 biological replicates for each condition were then used for comparisons (106,176 total) (Figure S4 and Table S2). A comparison of the two strains for each treatment condition using quartile normalized read counts demonstrated that the majority of promoter containing peaks (within 3kb of the TSS) and putative enhancer regions (>3kb from TSS) were consistent between the two strains across treatment conditions (Figure 3A-F).

**Figure 3.**
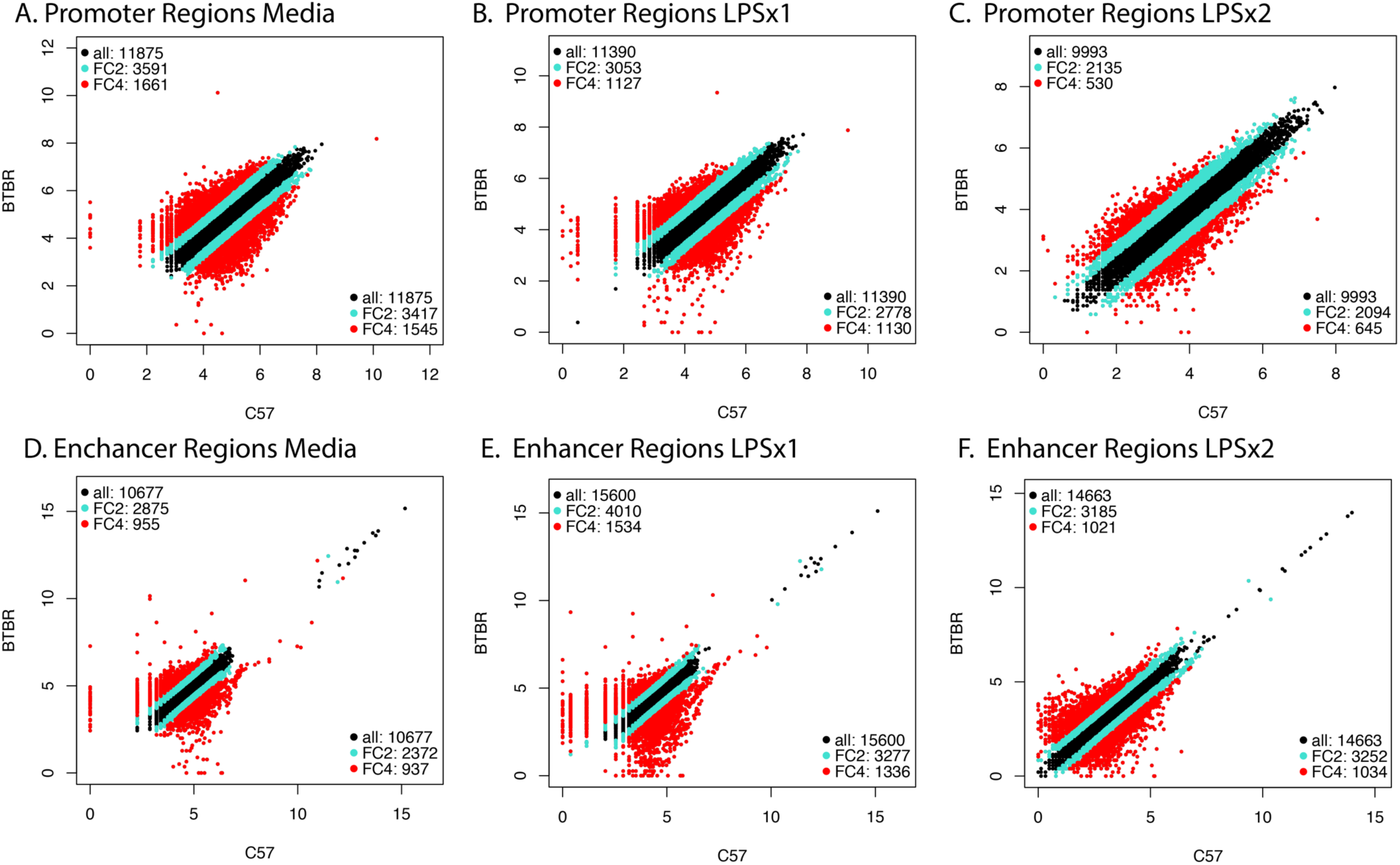
Correlation of quartile normalize ATAC-seq signal for all consensus peaks between strains. **A.** Promoter regions for C57 versus BTBR media peaks. **B.** Promoter regions for C57 versus BTBR LPSx1 peaks. **C.** Promoter regions for C57 versus BTBR LPSx1 peaks. **D.** Enhancer regions for C57 versus BTBR media peaks. **E.** Enhancer regions for C57 versus BTBR LPSx1 peaks. **F.** Enhancer regions for C57 versus BTBR LPSx2 peaks. For all panels, promoter regions were defined as within +/-3kb of the nearest gene TSS. Enhancer regions were all other non-promoter regions. Fold change (FC) > +/-2 is highlighted teal and FC >+/-4 is highlighted red.

To statistically test for regions with differential accessibility between strains and LPS treatments, we compared normalized read counts within the consensus peaks. Using a general linear model with effects for strain and LPS treatment and a correction for the within subject’s correlation from the repeated measures design, we identified 62,146 significant Differentially Accessible Regions (DARs) (Benjamini-Hochberg corrected *p*-value < 0.05) (Table S2 and Figure S5). Chromatin accessibility levels (normalized read counts within significant peaks) across samples clustered by both LPS condition and strain, highlighting impacts of both factors on chromatin accessibility (Figure 4A). A single LPS treatment resulted in a significant increase (adjusted *p*-value < 0.05) in chromatin accessibility in 4,764 peaks in C57 and 9,972 peaks in BTBR (LPSx1-DARs). A second LPS dose resulted in 6,499 peaks in C57 and 8,415 peaks in BTBR (LPSx2-DARs). Similarly, chromatin accessibility decreases were also observed in response to LPSx1 and LPSx2 (Figure 4B and C, Table S2) (adjusted *p*-value < 0.05).

**Figure 4.**
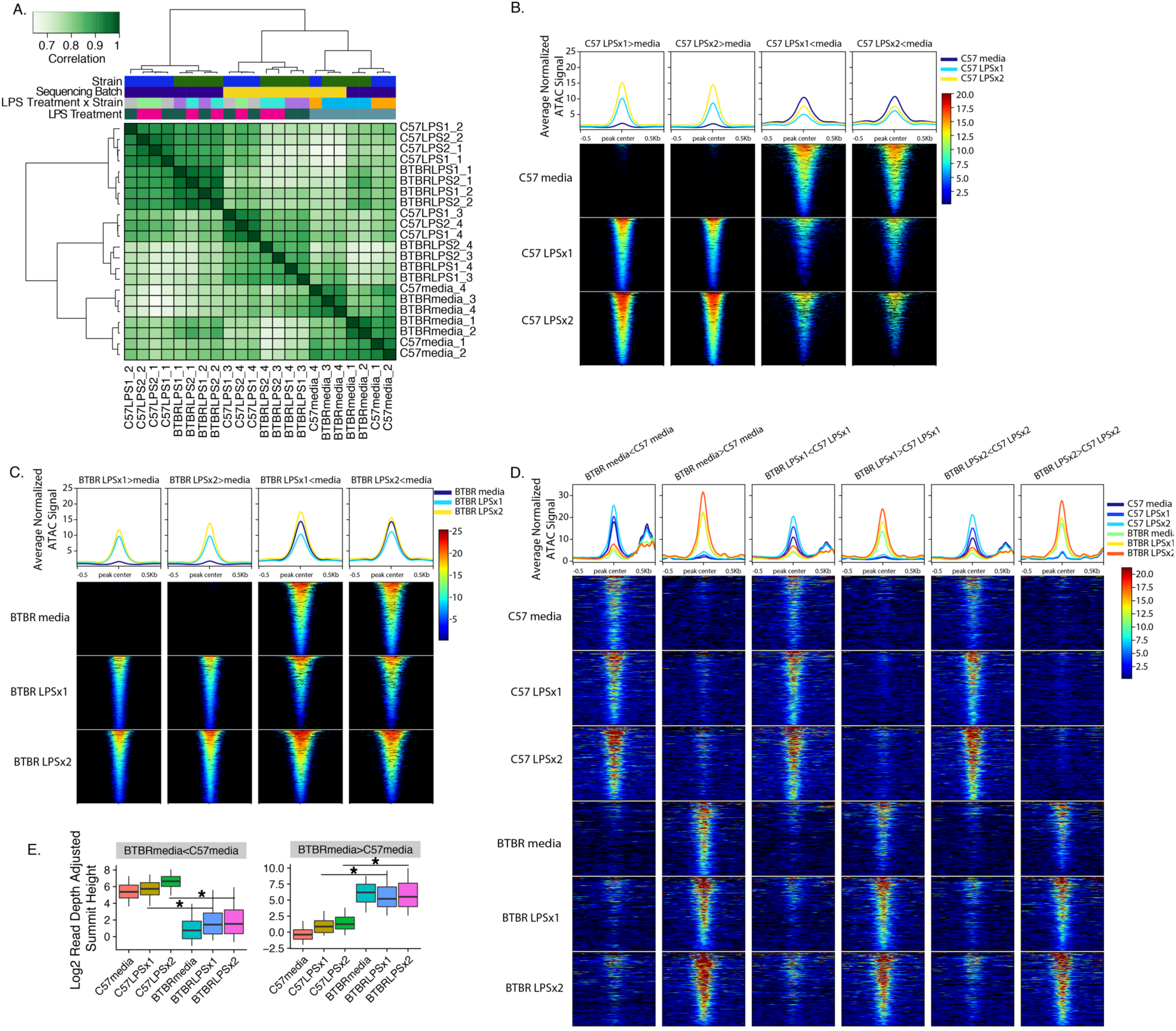
Strain-independent and Strain-specific Chromatin Responses to LPS. **A.** Clustered Heatmap of ATAC-seq signal (maximum read pileup value normalized to relative library size) for all significant peaks identified across all conditions. **B.** Average ATAC-seq signal (reads normalized to sequencing depth and averaged across replicates) for DARS identified in C57 in response to LPS. Line plots show means across DARs for each comparison. Each row of the heatmap shows signal averaged across individual DARs. **C.** Same as B but for BTBR. **D.** Same as B but for regions with differential accessibility between strains for each LPS treatment condition. **E.** Boxplots for the average ATAC-seq signal (reads normalized to sequencing depth) for DARs identified in the baseline condition. Boxes are defined by the 25^th^ and 75^th^ quartiles for each condition with median values shown as the horizontal line. Whiskers are defined by the 5^th^ and 95^th^ percentile values. ANOVA with Benjamini-Hochberg corrected posthoc comparisons \**p*-value < 0.05. All statistics and comparisons are shown in Table S5.

To compare the strains, we first examined the impact of strain on the baseline media condition, identifying 731 DARs (Figure 4C and Table S2). Peaks with lower accessibility in BTBR (367 BTBR media < C57 media) were significantly enriched within distal intergenic regions (Table S3) and showed enrichment (Fisher’s exact test) for previously published lists of enhancers sensitive to immune stimulation in BMDM cultures(Barish et al., 2010; Ghisletti et al., 2010; Link et al., 2018b) (immune enhancers, Figures S6 and Table S4). Peaks with higher accessibility in BTBR at baseline (364 BTBR media > C57 media) were also enriched in distal intergenic regions (Figure 4D, Table S3) but not in immune enhancers(Barish et al., 2010; Ghisletti et al., 2010; Link et al., 2018b) (Figure S6), suggesting that a small subset of immune-responsive enhancer elements are suppressed at baseline in the BTBR BDMD cultures.

Regions with strain-specific accessibility differences at baseline were also differential upon treatment with LPS (Figure 4D). Specifically, regions with lower accessibility in the BTBR media condition were less likely to be open upon LPS treatment compared to C57 and vice versa (Figure 4D). To directly test this observation, we compared the average normalized ATAC signal within the same regions identified by two sets of baseline peaks for all conditions (Figure 4E and Table S5). As predicted, regions showing lower accessibility in BTBR versus C57 at baseline also showed lower accessibility in response to LPS treatment (ANOVA, posthoc comparisons adjusted *p*-value < 0.05). In comparison, regions with higher accessibility in BTBR than C57 at baseline also showed higher levels of LPS response (ANOVA, posthoc comparisons adjusted *p*-value < 0.05) (Figure 4E, Table S5). Together, these results indicate that the baseline chromatin accessibility differences between the strains influence subsequent LPS response. Interestingly, the regions with lower accessibility in BTBR compared to C57 at baseline are also enriched for published immune regulatory enhancers(Barish et al., 2010; Ghisletti et al., 2010; Link et al., 2018b) (Figure S6), suggesting a difference in enhancer accessibility in the BTBR.

We next compared the response to LPS between the two strains. LPS-DARs identified in C57 were significantly shared with LPS-DARs identified in BTBR (Permutation overlap analysis FDR < 0.05) (Table S5). However, there were a small number of regions that showed differential accessibility in response to LPSx1 or LPSx2 between the strains (Figure 4D and Table S2). There were 633 peaks with BTBR>C57 in response to LPSx1 and 682 in response to LPSx2. There were 816 peaks with BTBR<C57 in response to LPSx1 and 826 in response to LPSx2. Regions with lower accessibility in BTBR were enriched for numerous immune regulatory enhancers(Barish et al., 2010; Ghisletti et al., 2010; Link et al., 2018b) (Figure S6), further supporting differential enhancer accessibility in response to LPS in BTBR.

### BTBR genetic variants are enriched within regions of differential chromatin accessibility between strains

To test the hypothesis that genetic differences between the two strains may influence their differential chromatin responsiveness, we utilized previously published BTBR genetic variants compared to C57 reference genome(Jones-Davis et al., 2013) (Figure 5). In BTBR, both SNPs and InDels (Insertions and Deletions) are distributed over numerous genomic features including splice sites, introns, UTRs, coding regions, promoters and intergenic regions (Figure 5A). To test the impact of BTBR genetic variants on chromatin accessibility, DARs were examined for enrichment of BTBR SNPs and InDels. Permutation testing compared each set of DARs to all consensus peaks. Strain-specific DARs at media baseline and in response to LPS were significantly enriched for both BTBR SNPs and InDels (Figure 5B and Table S6), while strain-independent LPS-DARs were not enriched for these genetic variants. These results support the hypothesis that natural genetic variation between strains may contribute to some of the altered chromatin state at baseline and in response to LPS.

**Figure 5.**
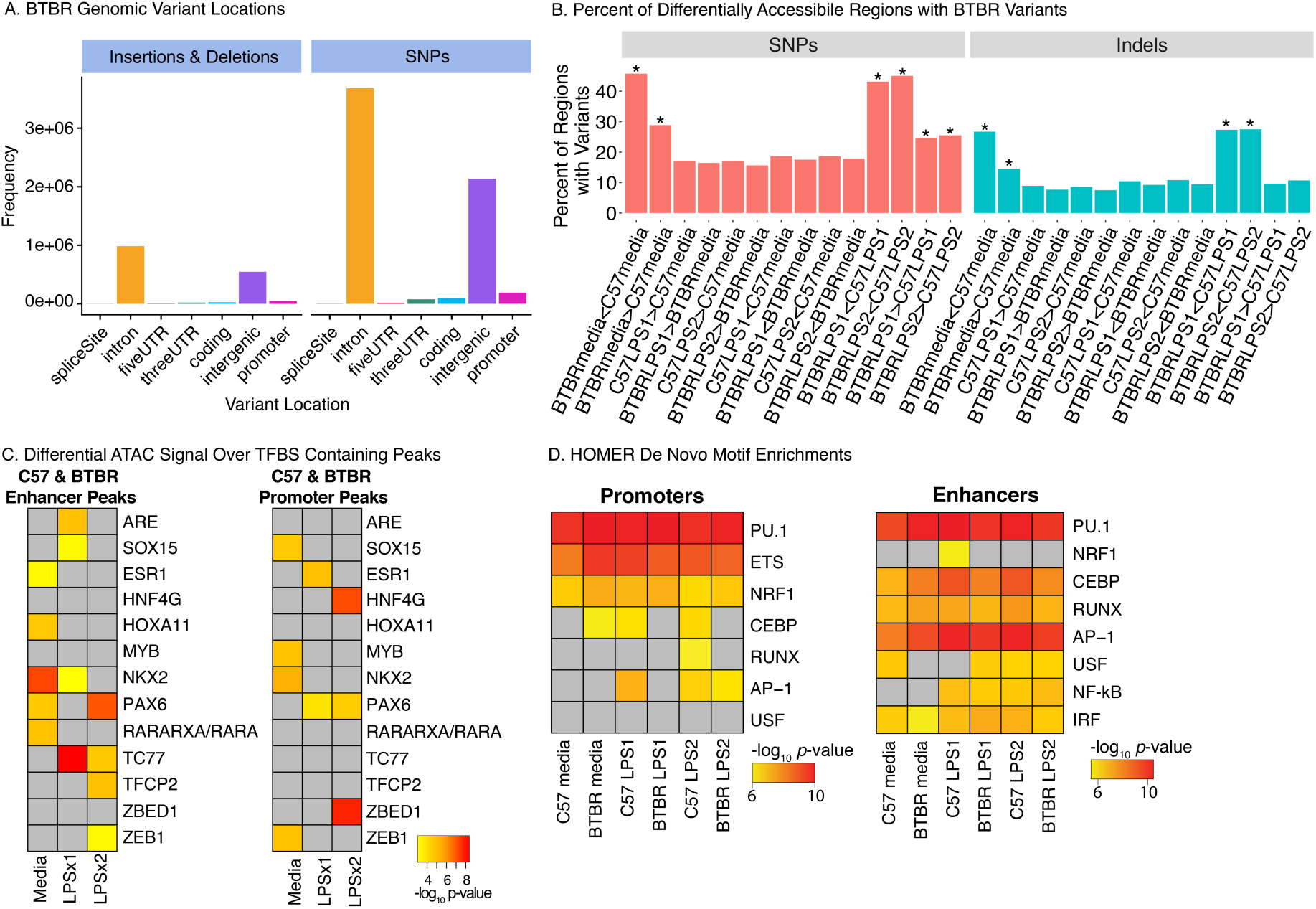
BTBR Genetic Variants Impact Chromatin Accessibility and Transcription Factor Binding Sites **A.** BTBR Insertions and Deletions (InDels) and SNPs from the Mouse Genomes Project (https://www.sanger.ac.uk/science/data/mouse-genomes-project) occur in numerous genomic elements including introns and intergenic regions. **B.** Percent of DARs containing BTBR SNPs or InDels. **\*** FDR q-value < 0.05 for permutation enrichment between BTBR variant and chromatin accessibility regions (see Table S6 for statistics). **C.** MMARGE analysis results showing chromatin accessibility between strains significantly differs for peaks containing these TF binding motifs with genetic variants. Peaks were defined for each treatment condition (consensus peaks) and subdivided into promoters (< 3 kb from TSS) and enhancers (non-promoters). **D.** *De novo* motif enrichment analysis was performed with HOMER for ATAC-seq consensus peaks of each strain and treatment condition. For C and D, significant enrichments are shown as -log_10_ *p*-values. Non-significant enrichments are in grey.

To further test this hypothesis that genetic variation between strains may alter accessibility to specific transcription factor binding motifs, variant analysis was performed for each peak set using MMARGE (Motif Mutation Analysis for Regulatory Genomic Elements)(Link et al., 2018c). MMARGE determines the significance of transcription factor (TF) binding motifs by evaluating correlation between the genetic variations interrupting the motif and the level of chromatin accessibility. MMARGE calculates the distribution of chromatin accessibility between two strains for all loci with the motif of interest in the first strain and compares it with the distribution of chromatin accessibility at all loci with the motif in the other strain. If the presence of the motif is important for chromatin accessibility, the distributions will not overlap and therefore be significant. We separated the loci into those within promoters (< 3 kb from TSS) compared to putative enhancers (non-promoters). Out of the 287 TF binding motifs tested, genetic variation leading to differences in TF binding motifs between the strains were significant for 13 different TF binding motifs (Figure 5C), suggesting that genetic variation between the strains may contribute to differential accessibility at these loci. For example, strain dependent chromatin accessibility was impacted by genetic variants at LPSx1 and LPSx2 responsive enhancers containing the motif of TCF7, a repressive transcription factor that plays important roles in the regulation of Runx splicing and has been implied in macrophage functions in various other tissues(Link et al., 2018a). In a complementary approach, we applied HOMER’s *de novo* motif analysis to the same set of consensus peaks divided into promoters and enhancers (Figure 5D). In enhancers and promoters across treatment conditions and strains, we found significant motif enrichments for the canonical macrophage transcription factors PU.1, CEBP, AP-1, but did not find evidence for genetic stain differences differentially impacting their binding motifs across conditions.

### Differential chromatin accessibility mirrors differential gene expression in response to LPS

To test the hypothesis that strain-specific differences in chromatin accessibility impact expression of tolerized and non-tolerized genes in response to LPS treatment, we compared DARs within promoters to DEGs from the array (Figure 6 and Table S7 and S8). We hypothesized that genes that respond to LPS with non-tolerized patterns of expression would open chromatin at their promoters in response to LPS and maintain that open state across multiple LPS treatments. As expected, DEGs showing a non-tolerized pattern of expression were significantly enriched for genes with promoter DAGs with increased accessibility upon LPSx1 and LPSx2 (Table S8). This was true across both strains, regardless of expression differences between strains. For example, the promoter region of *Socs3* is open and expression is maintained across LPS treatment conditions (Figure 6B) in both strains. This relationship suggests that promoters of non-tolerized genes in both strains maintain a permissive transcriptional state for on-going gene expression in response to repeated LPS.

**Figure 6.**
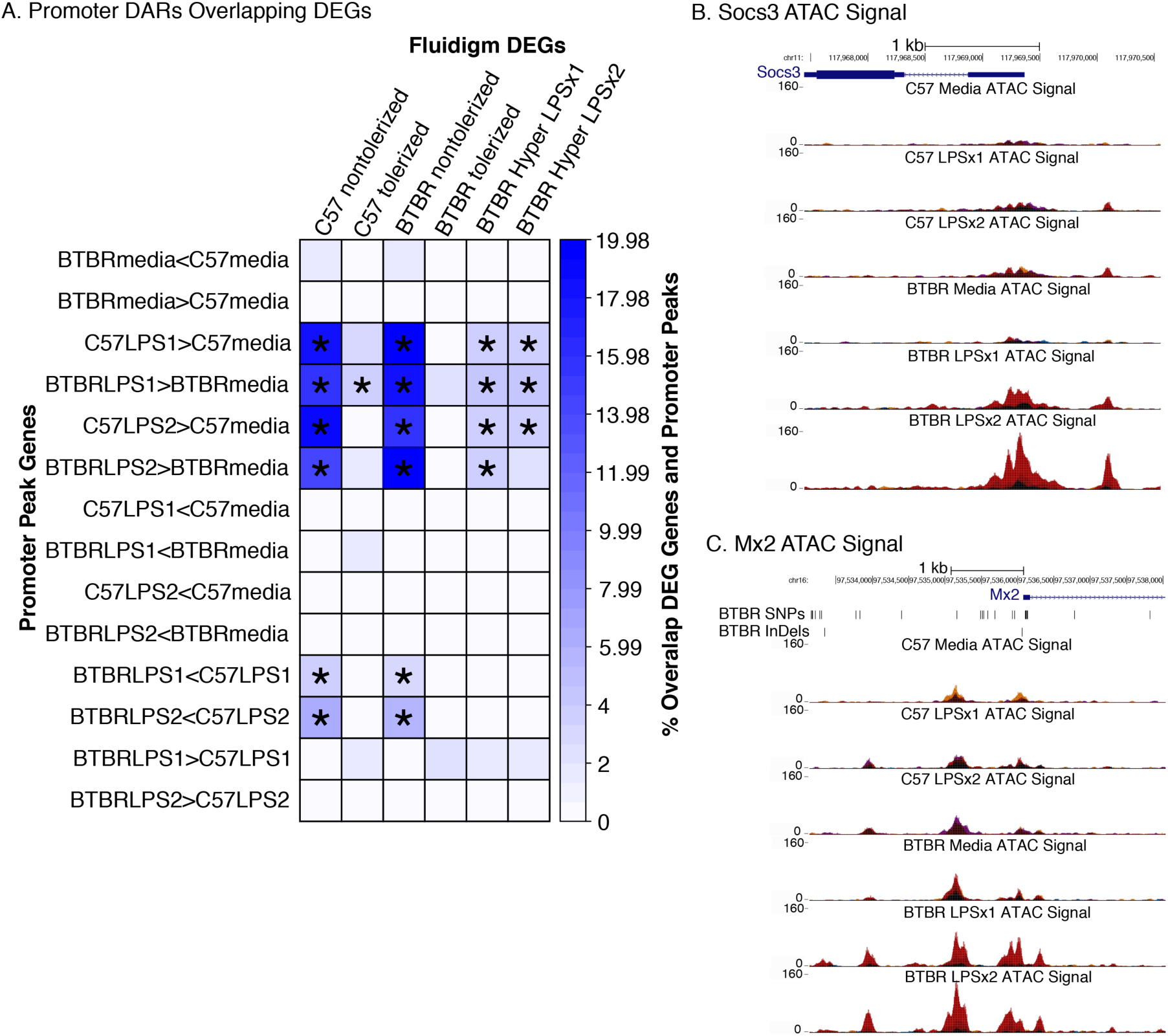
Strain-Specific Promoter DARs are Enriched for Non-Tolerized Genes **A.** Differentially expressed genes (DEGs) from Figure 2 were examined for enrichment of DARs within promoter regions of genes. The heatmap is colored by the percent of overlap between each row/column comparison. Significant enrichments (Fisher’s exact test) are denoted with * for al FDR *p*-value < 0.05. **B**. Normalized ATAC-seq signal over the promoter region of the *Socs3* gene. *Socs3* shows a non-tolerized pattern of gene expression (Figure 2B) and shows increased accessibility at its promoter across treatments. Replicates are shown in different colors for each UCSC genome browser track. **C**. Normalized ATAC-seq signal at the promoter region of the *Mx2* gene. *Mx2* shows a tolerized pattern of gene expression (Figure 2B) and hyper-responsive in expression of BTBR to LPS. Within each strain, the ATAC-seq signal at the *Mx2* promoter shows similar levels of accessibility across LPS treatments.

Similarly, we predicted that genes that become tolerized in response to repeated LPS doses would be enriched for peaks only present in response to LPSx1. However, we found a significant enrichment only for the BTBR LPSx1> media condition, but not for C57, suggesting that in C57 these genes are either already open at baseline (hence no difference between media and LPS treatment conditions) or regulated at a level other than simple chromatin accessibility. It also suggests that BTBR may have an exaggerated opening of chromatin in response to LPSx1 at tolerized gene promoters, potentially contributing to over-expression. For example, *Mx2* shows an open promoter chromatin state across treatment conditions, but a tolerized pattern of gene expression (Figure 6C). *Mx2* expression also shows hyper-responsive levels in BTBR for both LPS treatments which is paralleled by the increased accessibility (Figure 6C). Similarly, genes that hyper-responded in expression to LPS in the BTBR strain were also enriched for promoter DAGs that increased accessibility in response to LPS in BTBR and C57, supporting the conclusion that the transcriptional hyper-responsiveness may be related to the more open chromatin over promoters for these genes in BTBR.

## Discussion

This study relates naturally occurring genetic variation to changes in chromatin accessibility and gene expression in innate immune cells from a mouse model of ASD. Similar to previous findings in human monocytes(Enstrom et al., 2010), BTBR BMDM show a hyper-response to immune stimulation. We identified novel strain-specific genetic and epigenetic differences enriched within gene promoters showing enhanced expression in BTBR. This is the first work to directly examine how epigenetic differences regulate gene expression in BTBR immune cells and highlight the interaction between natural genetic variation, chromatin, and immune responsiveness. The alterations in BTBR immune function have been suggested to contribute to the development and manifestation of ASD relevant behaviors in BTBR(Heo et al., 2011; Onore et al., 2013). Several lines of evidence point to ongoing and prominent activation of innate myeloid immune cells in ASD(Ashwood et al., 2011; Enstrom et al., 2010; Hu et al., 2018; Inga Jácome et al., 2016; Molloy et al., 2006). Furthermore, we and others have shown increased activation of circulating myeloid cells in the periphery following *ex vivo* stimulation, including changes in gene expression and the release of pro-inflammatory cytokines(Ashwood et al., 2011; Enstrom et al., 2010; Mendizabal et al., 2016). The degree of myeloid cell activation was associated with more impaired behaviors and increased burden of co-morbidities in children with ASD(Ashwood et al., 2011). Our finding suggests that BTBR is a relevant model for studying the complex genetic architecture of immune alterations in ASD. Future work is needed to investigate natural human genetic variation in endotoxin tolerance responsiveness in human macrophages from patient populations.

We identified significant enrichments for BTBR strain variants within strain-specific regions of differential chromatin accessibility both under baseline conditions and in response to LPS, demonstrating an association between genetic variation and chromatin regulation of inflammatory response. These findings suggest that naturally occurring genetic variation in humans may also influence mechanisms for regulating innate immune responses and specifically responses to repeated pathogen exposure. Previous work in human monocytes has identified numerous genetic variants that contribute to regulation in response to immune stimulation(Fairfax et al., 2014) as well as in mediating the immune response during sepsis(Davenport et al., 2016). Understanding how individual genetic variation may impact chromatin regulation and gene expression in immune cells is critical for developing screening strategies for predicting individual patient outcomes and new treatment development(Davenport et al., 2016).

Natural genetic variation between mouse strains has also previously been shown impact both active and resting BMDM cultures(Heinz et al., 2013; Link et al., 2018a). In a comprehensive comparison of BMDM from 5 mouse strains (not including BTBR), strain-specific gene expression resulted primarily from genetic variation in distal regulatory elements(Link et al., 2018a). Under baseline conditions there was a large variation in gene expression with 10% of expressed transcripts varying by at least 4-fold between strains. In response to treatment with a Toll Like Receptor 4 agonist Kdo2-lipid A (KLA), distinctions between the strains emerged but the general response to KLA was conserved between strains, similar to our findings. The majority of these differences in gene expression were not explained by genetic differences in promoter sequences but instead by cis-variation in distal putative enhancers. There were even larger differences between strains in chromatin accessibility and binding of several transcription factors (TF), including Cebp, Pu.1, CJun, and P65 subunit of NfKB that were not explained by genetic strain differences alone. However, histone marks and long-range chromatin interactions revealed strain-dependent impacts of *cis*-regulatory domains on TF binding and gene expression(Link et al., 2018a). Our findings of BTBR versus C57 strain-specific differences in BMDM responsiveness are consistent with these earlier findings in identifying a highly complex set of genomic and epigenetic interactions that together cooperatively shape gene regulation in BMDM.

The relationship between chromatin accessibility and gene expression is complex, as chromatin state does not always directly predict expression(Link et al., 2018a). Prior work in BMDM in response to endotoxin tolerance identified loss in chromatin accessibility over the promoters of several exemplar tolerized genes(Foster et al., 2007). While this is consistent with our findings at some promoters, genes with tolerized expression in C57 were not generally enriched for LPS-DARs with decreased accessibility in C57. Similarly, not all genes with non-tolerized expression patterns in C57 showed more open promoters across LPS treatment conditions. Together this indicates the role of additional epigenetic and gene regulatory mechanisms that contribute to regulation of gene expression during the formation of endotoxin tolerance. Previous work has suggested critical roles for histone modifications(Foster et al., 2007; Novakovic et al., 2016; Saeed et al., 2014) as well as differential transcription factor binding in response to immune stimulation(Link et al., 2018a). Future work will be needed to more fully test the role of additional epigenomic mechanisms in regulating gene expression required for the formation of endotoxin tolerance in innate immune cells.

This work focused on immune gene regulatory mechanisms in bone marrow derived macrophages. BMDM cultures are a commonly used system for examining molecular mechanisms of immune function but may not fully capture all of the *in vivo* aspects of immune cell regulation. For example, the epigenome of tissue resident macrophages adapt to their tissue of residence providing a unique chromatin regulatory state dictating gene expression(Lavin et al., 2014). This may be extremely important in future investigation of tissue resident macrophages such as brain microglia that are exquisitely tuned to their local environments(Gosselin et al., 2017). Future work will be needed to characterize immune tolerance and chromatin regulation in additional tissue resident immune cell populations. Understanding the molecular mechanisms underlying the altered immune cell function in ASD models and human immune cell populations will lead to novel insights into immune dysfunction in ASD and identification of new immune-based therapeutic targets.

## Methods

### BMDM Cultures and LPS Treatment

All experiments were conducted in accordance with the National Institutes of Health Guidelines for the Care and Use of Laboratory Animals. All procedures were approved by the Institutional Animal Care and Use Committee of the University of California, Davis. Bone marrow cells were isolated from postnatal day 28 BTBR T^+^ *Itpr3*^*t*f^/J (Jax 002282) and C57BL6/J (Jax 000664) male mice. Isolated cells were cultured for seven days to generate bone marrow derived macrophages (BMDM) in DMEM/F12 media supplemented with 10% fetal bovine serum and 1% Penicillin-Streptomycin (base media). BMDM were then reseeded at a density of 5×10^5^ cells per well of a 6 well plate and incubated overnight. There were three treatment conditions: media, one dose LPS (LPSx1) or two doses of LPS (LPSx2). BMDM were treated with media (media or LPSx1) or 100ng LPS/ml for four hours (LPSx2). Cultures were then washed twice with HBSS and incubated overnight in base media. Twenty hours later cultures were treated with media (media) or 100ng LPS/ml for 24 hours (LPSx1 and LPSx2). Following treatment cells were collected for RNA or ATAC-seq.

### Fluidigm Biomark HD Delta Gene Expression Assay

Gene expression for selected tolerized and non-tolerized gene targets identified from (Foster et al., 2007) were examined in two independent experiments of LPS treatment of BMDM. Gene expression analysis was performed using the Fluidigm Biomark HD Delta gene assay on a 48.48 Integrated Fluidic Circuit. Delta gene assay design was performed by Fluidigm for equivalent amplification between the two genotypes and primer sequences are listed in Supplemental Table S2. An amount of 2.5ng of total RNA was used as input for cDNA synthesis (Fluidigm, 100-6472 B1) followed by 10 cycles of pre-amplification (Fluidigm, 100-5875 C1) using a mix of all forward and reverse primers. Samples were then diluted five-fold and analyzed on a 48.48 IFC and Biomark HD machine in duplicate or triplicate (Fluidigm, 100-9791 B1). Cycle threshold (Ct) values below detection were replaced with the max Ct value of 30. Ct values were averaged across technical replicates and then normalized to *Hprt* expression using the delta Ct method (Ct gene – Ct *Hprt*). The delta Ct values were then used for group comparisons by ANOVA with factors for strain, LPS treatment and the interaction with experimental batch as a covariate. For display purposes, the delta delta Ct values were then calculated relative to the average Ct value of the C57 media treated control for each gene and relative fold change expression levels were calculated for each group by 2^(-delta delta Ct).

Gene expression levels were defined as *non-tolerized* if expression increased significantly with one dose of LPS compared to media (corrected posthoc media vs LPSx1 *p*-value < 0.05) and there was not a significant difference between one and two doses of LPS (corrected posthoc LPSx1 vs LPSx2 *p*-value > 0.05). Genes were defined as *tolerized* if expression increased significantly with one dose of LPS compared to media (corrected posthoc media vs LPSx1 *p*-value < 0.05) and expression was lower with two doses of LPS compared to one dose of LPS (LPSx1>LPSx2 and corrected posthoc *p*-value < 0.05). Genes were defined as *sensitized* if expression increased significantly with one dose of LPS compared to media (corrected posthoc media vs LPSx1 *p*-value < 0.05) and expression was further increased with two doses of LPS compared to one dose of LPS (LPSx1<LPSx2 and corrected posthoc *p*-value < 0.05). Genes with a hyper induction in BTBR compared to C57 were categorized for *Hyper LPSx1* (corrected posthoc C57 LPSx1 vs BTBR LPSx1 *p*-value < 0.05 and C57 LPSx1 < BTBR LPSx1) and *Hyper LPSx2* (corrected posthoc C57 LPSx2 vs BTBR LPSx2 *p*-value < 0.05 and C57 LPSx2 < BTBR LPSx2). *Media Baseline* differences were defined by a significant corrected posthoc comparison between C57 and BTBR media conditions with a *p*-value < 0.05.

### ATAC-seq Library Preparation

ATAC-seq was performed as described in(Buenrostro et al., 2015). Briefly, cultures of 50,000 BMDM were washed once in ice cold 1xPBS and then resuspended gently in 100µl ATAC-seq lysis buffer (1 mM TrisCl, pH 7.4, 1 mM NaCl, 0.3 mM MgCl2, and 0.01% Triton-X 100) to release nuclei. Nuclei were then pelleted (1,000xg, 10min, 4°C) and transposed in 2.5µl Tn5 (Nextera Tn5 transposase, Illumina), 12.5µl TD reaction buffer and 10.5µl water for 40min at 37°C with rotation. Following transposition, DNA was purified (MinElute PCR Purification Kit, Qiagen) and stored at −80°C. Samples were then thawed on ice and unique barcodes(Buenrostro et al., 2015) were added to each sample using 5 cycles of PCR(Buenrostro et al., 2015). A side PCR reaction was taken and used as input for qPCR for an additional 20 cycles to determine the optimal number of additional PCR cycles for each library (the cycle number that corresponds to one-third of the maximum fluorescent intensity). Each sample was then amplified by the determined number of PCR cycles and then purified (MinElute PCR Purification Kit, Qiagen). Final libraries were then size selected for fragments between 100-1000bp using AMPure XP beds (Beckman Coulter) and quality was checked by DNA High Sensitivity bioanalyzer. Libraries were pooled (6 per pool) and sequenced across four lanes of the HiSeq2500 to generate 50bp paired end reads.

### BTBR genome analysis

BTBR SNPs and Indels were downloaded from the Mouse Genomes Project (https://www.sanger.ac.uk/science/data/mouse-genomes-project) and transferred from mm9 to mm10 using CrossMap (version 0.2.8)(Zhao et al., 2014). The program MMARGE (Motif Mutation Analysis for Genomic Elements, version 1)(Link et al., 2018c) was used to create a modified version of the mm10 genome that incorporated BTBR SNPs and Indels (MMARGE.pl prepare_files). The BTBR genome was used to generate bowtie2 index files for subsequent alignments of BTBR sequencing data. SNPs and InDels were examined for variant location using the locateVariants function in the R package VariantAnnotation(Obenchain et al., 2014) and the mouse organism database (org.Mm.eg.db).

### ATAC-seq Alignment, Quality Control, and peak calling

All fastq files were trimmed using trimmomatic (version 0.38) to remove paired end adapters, minimum MAPQ score of 30, leading and trailing 3bp, and to a minimum length of 15bp. Trimmed fastq files were then aligned to either the mm10 genome or a modified mm10 that incorporated BTBR genetic changes using bowtie2(Langmead and Salzberg, 2012) with MMARGE(Link et al., 2018c) with default settings for paired end data. BTBR aligned data were then shifted back to mm10 standard genome coordinates using MMARGE.pl shift(Link et al., 2018c). For all samples aligned reads were then filtered using samtools (version 1.8) to remove PCR duplicates, unpaired reads, unmapped reads, and secondary alignments. To assess transposition efficiency and the proportion of nucleosome free reads compared to mono and di-nucleosomes, histograms of the distribution of insert sizes was created using picard-tools (version 2.18.4) CollectInsertSizeMetrics. The filtered reads were then shifted to account for the Tn5 insertions site (+4bp to positive strand and −5bp from negative strand) using the perl script ATAC_BAM_shifter_gappedAlign.pl (https://github.com/TheJacksonLaboratory/ATAC-seq/blob/master/auyar/). Shifted reads were filtered for nucleosome free regions (read length < 147bp) and mitochondrial reads were removed using samtools. Biological replicates for each strain and treatment condition were compared for consistency using deeptools(Ramírez et al., 2016). For each sample, bamCoverage was used to make bigwig files with read coverage normalized to 1x sequencing depth using an effective genome size for mm10 of 2,652,783,500 with chrM, chrX and chrY excluded from normalization. The bin size was set to 10bp and the mm10 blacklist regions were excluded. Biological replicates were then compared within each condition using multiBigwigSummary followed by plotCorrelation with --corMethod spearman –skipZeros.

Filtered reads were then used for peak calling with HOMER(Heinz et al., 2010) (-style factor -center -F 0 -L 0 -C 0 -fdr 0.01 -minDist 200 -size 200). The resulting peak files were then filtered to remove mm10 blacklist regions using bedtools. Biological replicates were compared using a Spearman’s correlation with Deeptools(Ramírez et al., 2016). Two C57 samples failed initial QC and were removed from subsequent analysis (see results). Alignments, read counts, filtering and peak counts were examined for differences between experimental conditions using a repeated measures ANOVA with factors for treatment, genotype and the interaction as well as a repeated measures factor for subject.

### ATAC-seq Differential Accessibility Analysis

Differential accessibility regions were identified from HOMER peaks called in NFRs using the R packages DiffBind(Stark and Brown, 2011), edgeR(Robinson et al., 2010), limma and voom(Ritchie et al., 2015). Individual peak files for each sample were combined into a dba object and the configuration was set for paired end sequencing. Consensus peaks between biological replicates for each condition were defined using dba.peakset as peaks occurring within 3 out of 4 biological replicates with peak overlap of at least 1bp. The fraction of reads in peaks (FRIP) was then calculated for each sample for all consensus peaks as an additional QC measure. Raw reads were then counted within all consensus peaks for each sample with summits set to True, and scaling to control samples set to false using Diffbind dba.count. The resulting counts were normalized for the total sequencing library size (after filtering described above) using edgeR with calcNormFactors and method set to Relative Log Expression (RLE), where the median library is calculated from the geometric mean of all columns and the median ratio of each sample to the median library is taken as the scale factor(Robinson et al., 2010). The RLE normalized counts were then further normalized for common, trended and tagwise dispersion using a design matrix with factors for genotype and treatment and a zero intercept. To account for the repeated measures aspect of the experimental design, the dgList object was then passed to voom and sample weights were calculated using the duplicateCorrelation function with block set to sample replicate. The final call to lmFit included the blocking factor and the correlation, resulting in a model that accounts for the correlation within subjects. Individual planned comparisons were then conducted using makeContrasts, contrast.fit, eBayes, and topTable. All significant regions (Benjamini-Hochberg corrected p < 0.05) were then further separated by the direction of accessibility change between conditions.

To examine the overlap between region conditions, region files were compared using the resampleRegions function for the number of overlaps in the R package regioneR(Gel et al., 2016). Region lists were tested for either more or less overlap than expected by chance using 1000 permutation tests of random resampling from the universe of all consensus peaks identified in the experiment. The resulting permuted *p*-values were then FDR corrected. The same approach was used to overlap regions with BTBR SNPs and InDels. Pairwise overlap heatmaps were made using the R package Intervene(Khan and Mathelier, 2017).

For overlapping regions with DEGs from the Fluidigm array, regions were annotated using the R package ChIPseeker with the overlap function set to “all”, which sets the gene overlap with the peak to be reported as nearest gene independent of whether it is a TSS region or not.

### Visualization of Average ATAC-seq Signal Over Peak Regions

Visualization of average ATAC-seq signal was done using Deeptools(Ramírez et al., 2016). Average bigwig files for each condition were made using (http://wresch.github.io/2014/01/31/merge-bigwig-files.html) from the read depth normalized bigwig files described above. The resulting bigwig files were then compared to specified peak regions using Deeptools(Ramírez et al., 2016) computeMatrix reference-point --referencePoint center -b 500 -a 500 –skipZeros followed by plotHeatmap --refPointLabel “peak center” --averageTypeSummaryPlot mean -- perGroup --colorMap RdBu --yMax 40.

### Region Enrichments

Enrichment of ATAC-seq regions within previously published regulatory regions was performed using a the R package LOLA(Sheffield and Bock, 2015) with a one-tailed Fisher’s exact test with FDR correction to p<0.05 across all comparisons. Region lists include regions identified in ENCODE ChIP-seq for mouse BMDM (www.encodeproject.org), and ChIP-seq target regions from several transcription factors in mouse BMDM with and without immune stimulation(Barish et al., 2010; Ghisletti et al., 2010; Link et al., 2018b).

### MMARGE’s mutation analysis

Consensus bigwig file from each condition were merged between the different strains with HOMER’s mergePeaks and split into promoter and enhancer regions based on their distance to the closest TSS (peaks less than 3kb from the next TSS were defined promoters, the rest enhancers). These files were annotated with raw read counts using HOMER and furthermore quantile normalized using R’s normalization. MMARGE’s mutation_analysis(Link et al., 2018c) was performed on each peak file using natural genetic variation from C57 and BTBR, as well as -no_correction. The results of this analysis were summarized with MMARGE’s summary_heatmap and significance threshold of 0.05.

### Modified Homer Motif Enrichment Analysis

To analyze enriched *de novo* motifs, MMARGE’s modified *de novo* motif analysis from HOMER was used. The analysis was performed on each peak file from each treatment in each strain for promoters, as well as enhancers. The original sequence per peak was extracted and HOMER’s de novo motif finding algorithm was used for these sequences.

### Scatterplots

To visualize the ATAC-seq data in scatter plots, peak files generated for MMARGE’s mutation analysis were used and plotted in R as log2 values.

## Acknowledgements

This work was supported by the National Institutes of Health (R01ES021707, R01NS081913 to J.M.L., RO1HD090214, RO1 MH118209, R21MH116383 to P.A.), Johnson Foundation. Brain & Behavior Research Foundation (NARSAD Young Investigator Award to A.V.C.), National Institutes of Mental Health (1K01MH116389-01A1 K01 Mentored Research Scientist Development award to A.V.C). This work used the Vincent J. Coates Genomics Sequencing Laboratory at UC Berkeley (NIH S10 OD018174 Instrumentation Grant) and the University of California, Davis Intellectual and Developmental Disabilities Research Center (IDDRC) (NIH U54 HD079125)

## Author Contributions

A.V.C., M.C., P.A. and J.M.L. designed the experiments. M.C. performed cell isolations, cultures, and LPS treatments. A.V.C. made ATAC-seq libraries and performed the gene expression experiments. A.V.C. and V.M.L. performed all data analysis. A.V.C., V.M.L., P.A. and J.M.L contributed to the manuscript writing.

