## Supplemental figures and legends for "Genetic variants drive altered epigenetic regulation of endotoxin tolerance in BTBR macrophages"

### Supplemental Figures and Table Legends

A. Insert Size Distribution

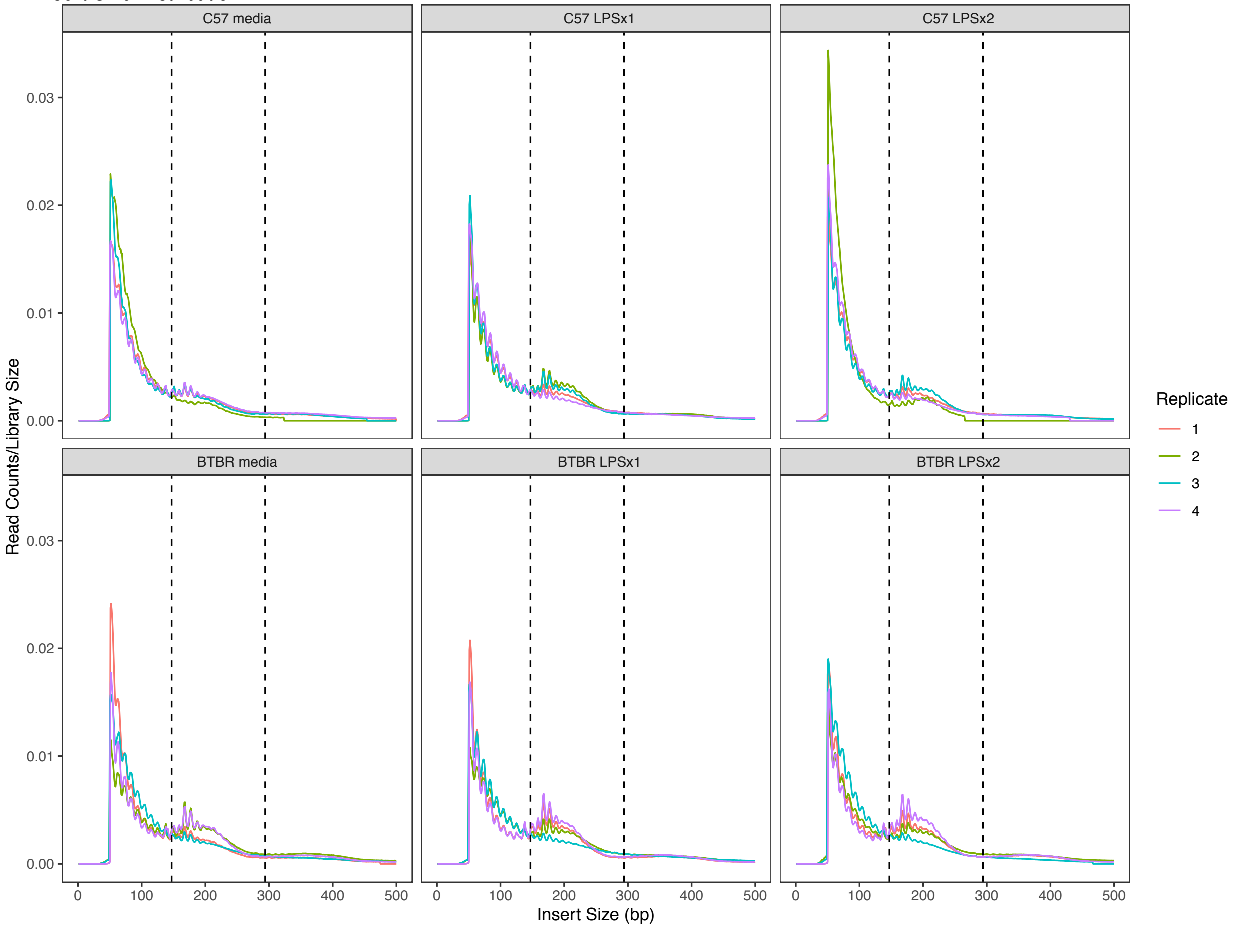


**FigureS1. ATAC-seq Library Scaled Insert Size Distribution**

ATAC-seq library insert size distributions for each biological replicate for each condition as assessed by Picard tools CollectInsertSizeMetrics. Size distributions are scaled as a ratio relative to the total library size to give Read Counts/Library. Dotted lines mark the nucleosome (147bp) and di-nucleosome (249bp) regions.

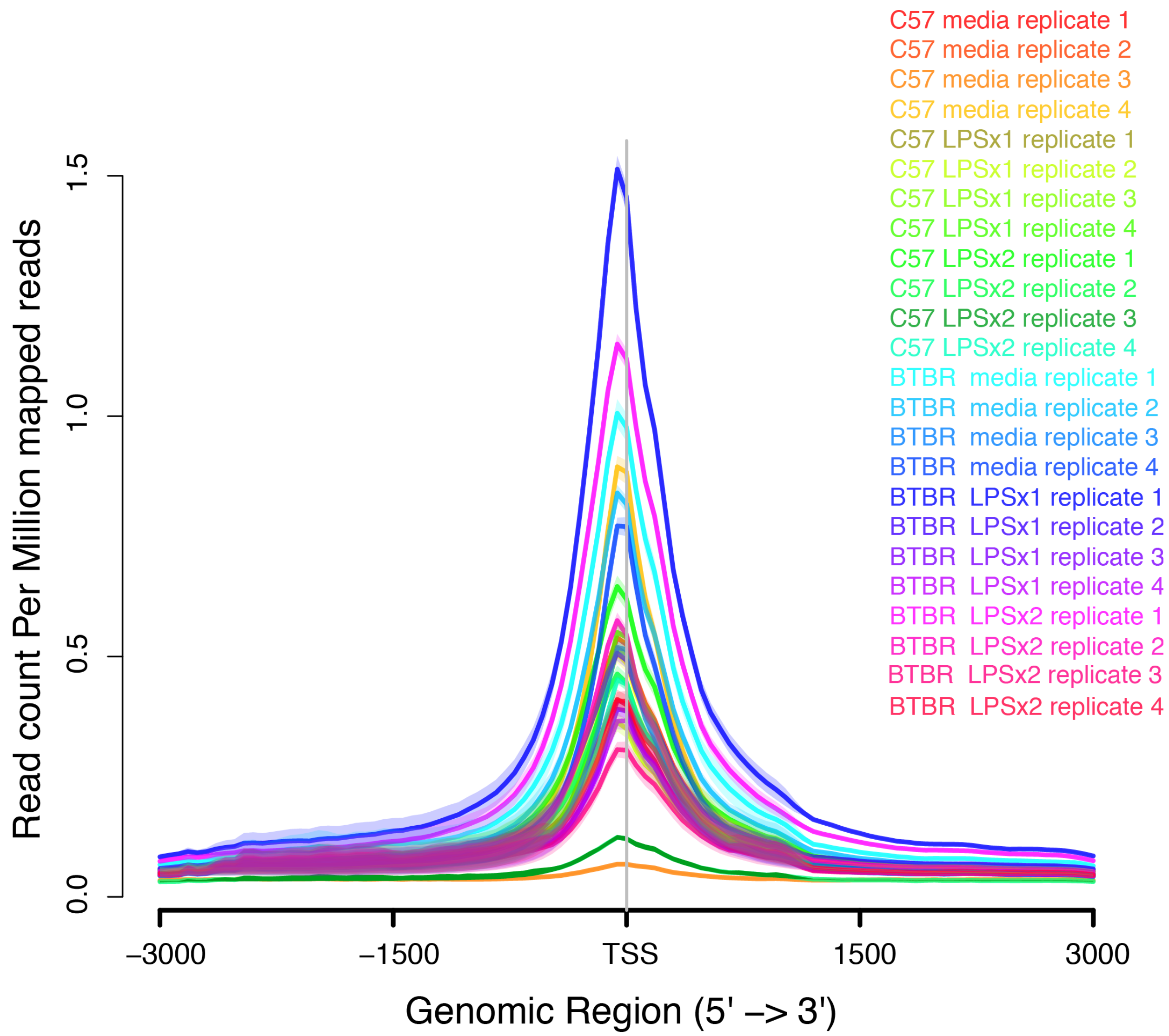


**FigureS2. Average ATAC-signal over all Transcription Start Sites (TSS).**
The Read Depth normalized ATAC-signal was visualized over all mouse (mm10) TSS to assess general quality of each ATAC library. Note the two samples with nearly flat lines (C57 media replicate 3 and C57 LPSx2 replicate 3) that were removed from the final analysis for also failing additional QC metrics.

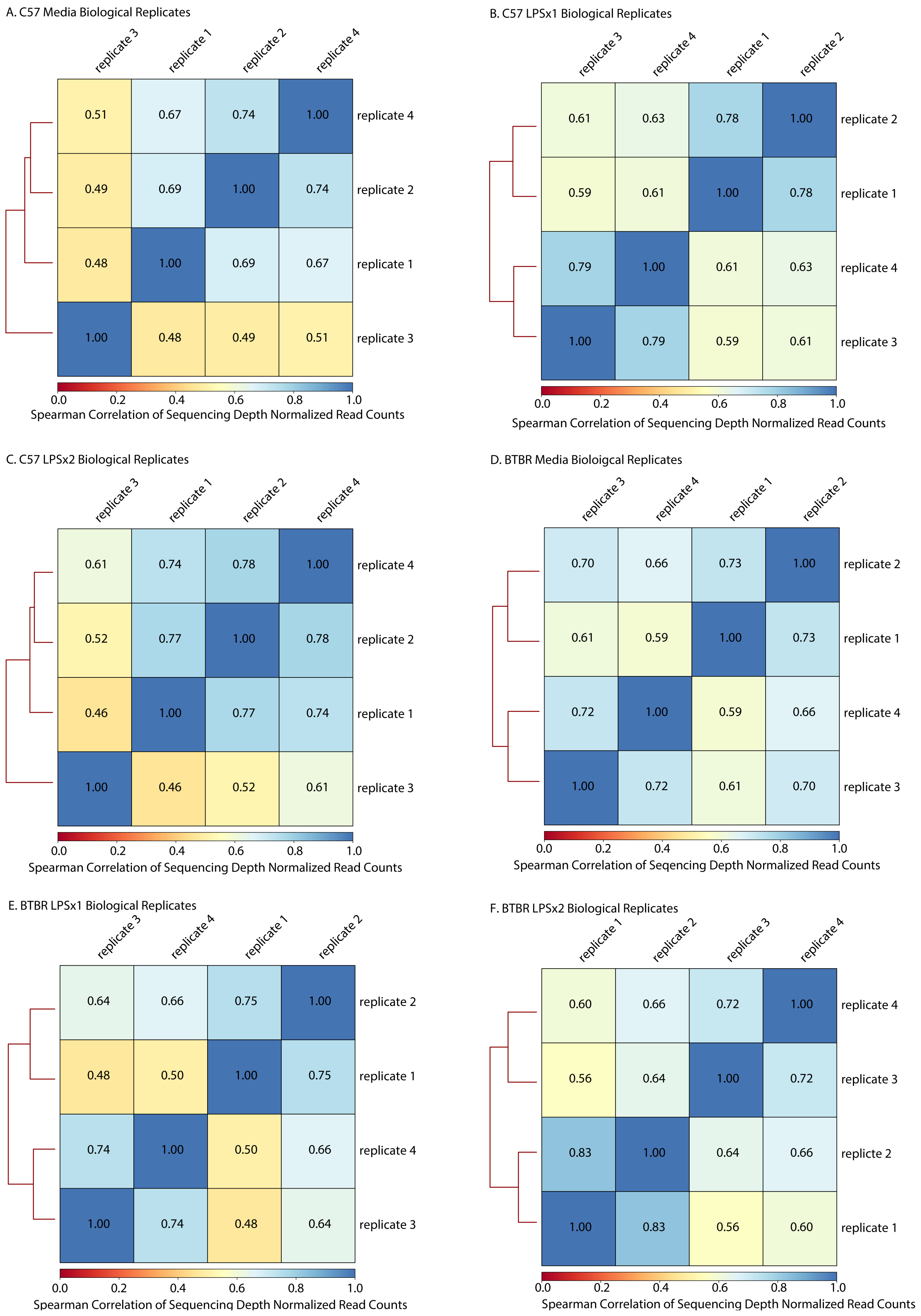
A. C57 Media Biological Replicates
replicate 3
replicate 1
replicate 2
replicate 4
0.51 0.67 0.74 1.00 replicate 4
0.49 0.69 1.00 0.74 replicate 2
0.48 1.00 0.69 0.67 replicate 1
1.00 0.48 0.49 0.51 replicate 3
0.0 0.2 0.4 0.6 0.8 1.0
Spearman Correlation of Sequencing Depth Normalized Read Counts
B. C57 LPSx1 Biological Replicates
replicate 3
replicate 4
replicate 1
replicate 2
0.61 0.63 0.78 1.00 replicate 2
0.59 0.61 1.00 0.78 replicate 1
0.79 1.00 0.61 0.63 replicate 4
1.00 0.79 0.59 0.61 replicate 3
0.0 0.2 0.4 0.6 0.8 1.0
Spearman Correlation of Sequencing Depth Normalized Read Counts
C. C57 LPSx2 Biological Replicates
replicate 3
replicate 1
replicate 2
replicate 4
0.61 0.74 0.78 1.00 replicate 4
0.52 0.77 1.00 0.78 replicate 2
0.46 1.00 0.77 0.74 replicate 1
1.00 0.46 0.52 0.61 replicate 3
0.0 0.2 0.4 0.6 0.8 1.0
Spearman Correlation of Sequencing Depth Normalized Read Counts
D. BTBR Media Bioloigcal Replicates
replicate 3
replicate 4
replicate 1
replicate 2
0.70 0.66 0.73 1.00 replicate 2
0.61 0.59 1.00 0.73 replicate 1
0.72 1.00 0.59 0.66 replicate 4
1.00 0.72 0.61 0.70 replicate 3
0.0 0.2 0.4 0.6 0.8 1.0
Spearman Correlation of Seqencing Depth Normalized Read Counts
E. BTBR LPSx1 Biological Replicates
replicate 3
replicate 4
replicate 1
replicate 2
0.64 0.66 0.75 1.00 replicate 2
0.48 0.50 1.00 0.75 replicate 1
0.74 1.00 0.50 0.66 replicate 4
1.00 0.74 0.48 0.64 replicate 3
0.0 0.2 0.4 0.6 0.8 1.0
Spearman Correlation of Sequencing Depth Normalized Read Counts
F. BTBR LPSx2 Biological Replicates
replicate 1
replicate 2
replicate 3
replicate 4
0.60 0.66 0.72 1.00 replicate 4
0.56 0.64 1.00 0.72 replicate 3
0.83 1.00 0.64 0.66 replicte 2
1.00 0.83 0.56 0.60 replicate 1
0.0 0.2 0.4 0.6 0.8 1.0
Spearman Correlation of Sequencing Depth Normalized Read Counts

**FigureS3. Spearman's Correlations Between Biological Replicates**

The Read Depth normalized ATAC-signal was compared between biological replicates within each strain x treatment condition using Deeptools bamCoverage. Chromosome M, X and Y and all mm10 blacklist regions were excluded from the analysis. Note the two samples with predominantly low correlations (C57media replicate 3 and C57LPSx2 replicate 3) that were removed from the final analysis. **A.** Spearman's Correlation coefficients for C57 media replicates. **B.** Spearman's Correlation coefficients for C57 LPSx1 replicates. **C.** Spearman's Correlation coefficients for C57 LPSx2 replicates. **D.** Spearman's Correlation coefficients for BTBR media replicates. **E.** Spearman's Correlation coefficients for BTBR LPSx1 replicates. **F.** Spearman's Correlation coefficients for BTBR LPSx2 replicates.

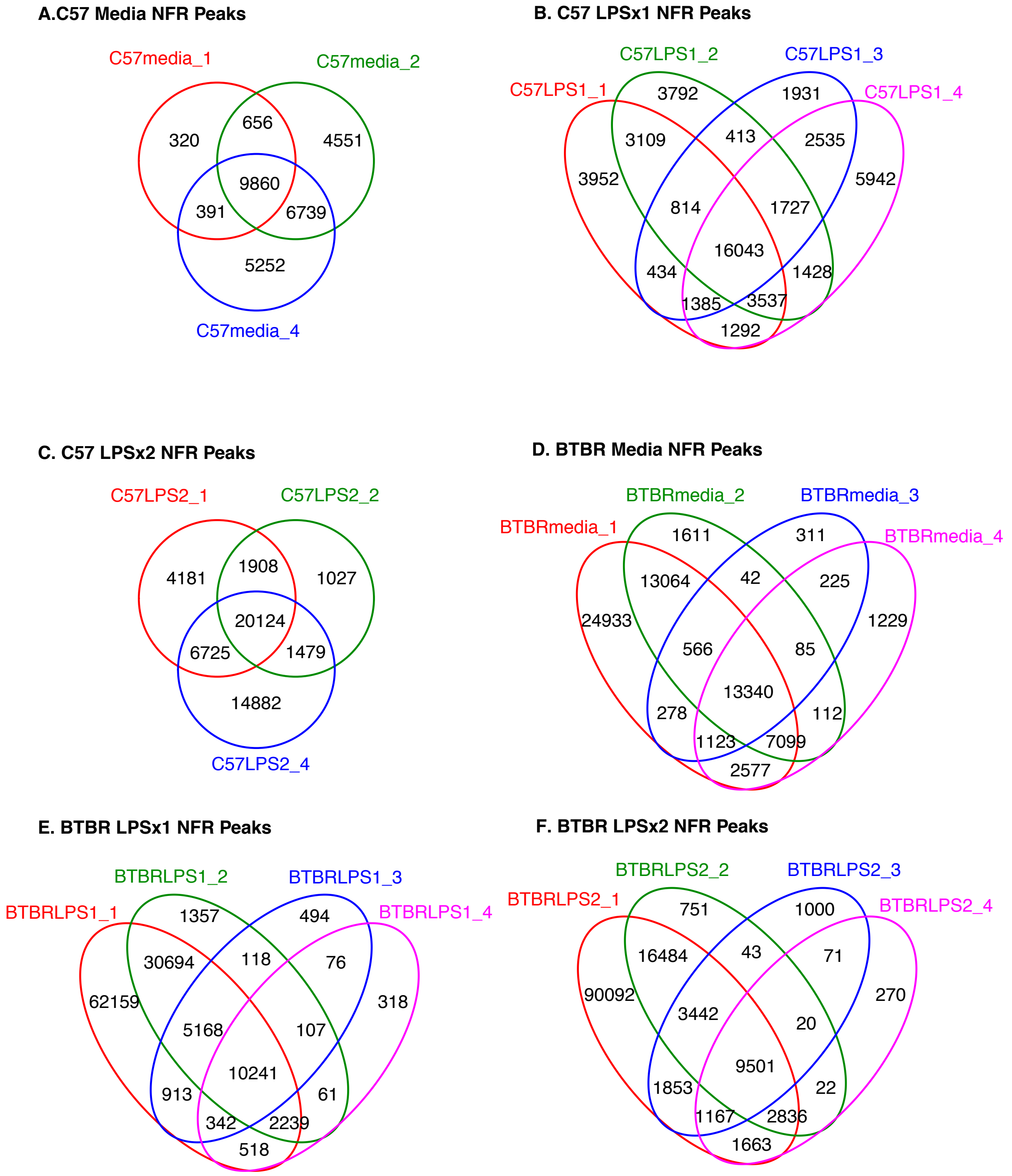


**FigureS4. Consensus Regions Across Strain and LPS Treatments**
**A.** HOMER peaks within Nucleosome Free Regions (NFR) for each biological replicate for the C57 BMDM media condition. Consensus peaks were defined as peaks occurring in at least 3 out of 4 replicates. **B.** Same as A. but for C57 BMDM LPSx1. **C.** Same as A. but for C57 BMDM LPSx2. **D.** Same as A. but for BTBR BMDM media. **E.** Same as A. but for BTBR BMDM LPSx1. **F.** Same as A. but for BTBR BMDM LPSx2.

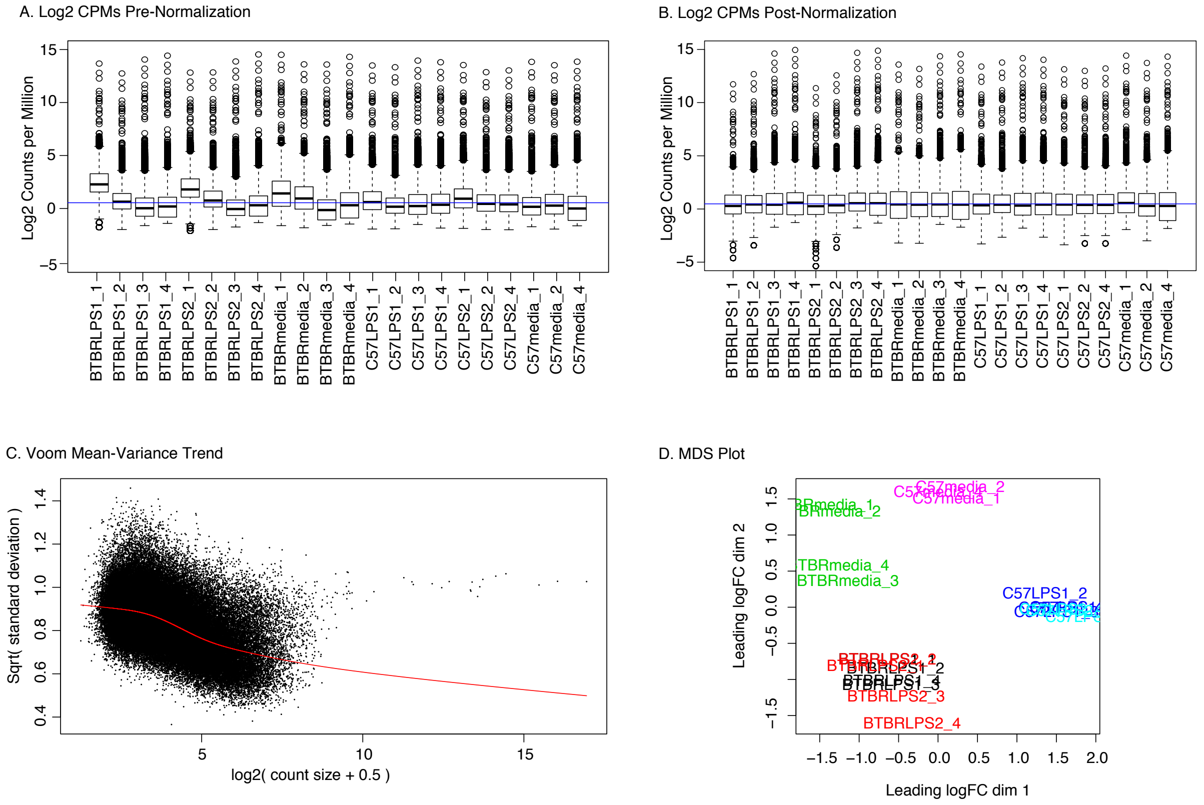


**Figure S5. Differential ATAC-Peak Analysis**

**A.** Log2 Counts Per Million reads sequenced for all consensus peaks identified across all conditions prior to any normalization. **B.** Log2 Counts Per Million reads sequenced for all consensus peaks identified across all conditions after Relative Log Expression (RLE) normalization, normalized for common, trended and tagwise dispersion using a design matrix with factors for genotype and treatment and a zero intercept. **C.** Mean-Variance relationship and trendline from limma-voom using a design matrix with factors for genotype and treatment and a zero intercept and the duplicateCorrelation function with block set to sample replicate. **D.** Multidimensional scaling (MDS) plot of all samples, colored by condition.

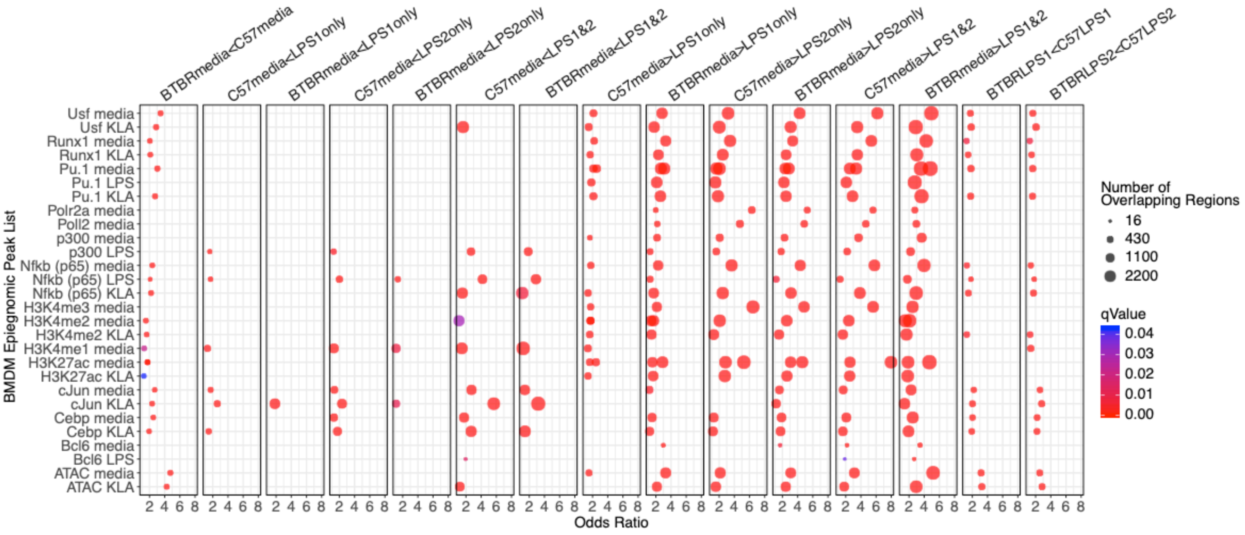


**Figure S6. Significant Overlaps Between DARs and Published BMDM Epigenomic Datasets**

**A.** Significant enrichments between identified differential strain and LPS treatment regions (DARs) with previously published epigenomic datasets in BMDM under baseline media conditions or in response to either LPS or Kdo2 lipid A (KLA), a TL4 agonist (Barish et al. 2010; Ghisletti et al. 2010; Link et al. 2018). Significant overlaps (Fisher's exact test $q<0.05$) are shown for each comparison as a dot. Dot size is scaled to the number of overlapping regions and colored by q-value. Cases with multiple dots per comparison list indicate significant overlap with more than one published dataset for that epigenomic mark.

**Supplemental Table Legends**

**TableS1. Fluidigm qRT-PCR Array Gene Expression Analysis**
Fluidigm Primers: Primer information for Fluidgim custom qPCR array design for genes from [3]. Class NT are non-tolerized genes, Class T are tolerized genes. Three additional potential housekeeping genes were included as well as gene targets previously shown to be misregulated in BTBR mice[4]. Primers were designed to amplify equivalently across both strained based on BTBR SNP information.
Fluidigm Samples: The number of biological replicates per condition for each target gene on the qPCR array.
Fluidigm ANOVA: ANOVA output results for each gene for delta Ct values from the qPCR array. The statistical model used was: lme(deltaCt ~ Strain * treatment + batch, ~1|sample).
Fluidigm Results: Average Fold Change values for each condition and gene, categorization (see methods) based on expression levels and Tukey HSD corrected posthoc comparisons. Relevant posthocs are included for columns beginning with adj.pval. MediaDifferences for are based on a significant (baseline differences) or non-significant (no baseline differences) posthoc comparison between C57 media and BTBR media conditions. The final Category column is based on whether the gene is tolerized, nontolerized or sensitized.

**Table S2. QC Data BMDM ATAC-seq**
SampleInfo: Information on each sample replicate including strain, treatment, sequencing batch, additional PCR cycles for library preparation, HiSeq2500 lane, and Sample Index.
ATACseqQC: For each sample read numbers are given for each step of the analysis including pre and post filtering, the number peaks in NFR called for HOMER, and the fraction of reads in peaks (FRIP) for each sample replicate is given. Samples with NA values for peaks and FRIP were excluded from the analysis for failing prior QC measures.
RMANOVA_ATACQC: Repeated measures ANOVA for each QC metric using a statical model for strain, treatment and the interaction (Strain:Treatment).
Consensus Peak Count: The number of consensus peaks for each sample and condition (in ¾ replicates per condition).
Differential Peak Count: The number of peaks per condition for each comparison. For each strain the total proportion of each set of peaks is calculated relative to all LPS responsive peaks for that strain.

**Table S3. Fisher's Exact Enrichment for Genomic Elements**
Fisher's Exact Test: One tailed Fisher's Exact Test for enrichment of each peakset within different genomic elements. All comparisons were then FDR corrected. Annotation was performed with ChIPSeeker.

**Table S4. Fisher's Exact Enrichment for BMDM Epigenomic Datasets**
Fisher's Exact Test: One tailed Fisher's Exact Test for enrichment of each peak set with previously published epigenomic datasets in BMDM under baseline media conditions or in response to either LPS or Kdo2 lipid A (KLA), a TL4 agonist (Barish et al. 2010; Ghisletti et al. 2010; Link et al. 2018). All enrichments were FDR corrected ($q<0.05$).

**Table S5. EnrichPeakOverlaps**
RegioneR Overlaps: Permutation testing for overlap enrichment between DAR peak lists by resampling against a random null distribution selected from all consensus peaks using the RegioneR package in R. All permutations were FDR corrected to q<0.05.
RM ANOVA peak signal: Results from repeated measures ANOVAs for the average ATAC signal within media peak sets across conditions. The statistical model included factors for strain, LPS treatment and the interaction with a random effect of biological replicate.
FDR Posthocs peak signal: FDR corrected posthoc tests for C57 versus BTBR for LPSx1 and LPSx2 conditions following the RM-ANOVA analysis of peak signal over each peak set.

**TableS6. BTBR Genomic Analysis**
BTBR variant summary: Counts of the number of BTBR variants in different genomic elements
BTBR Variant - Region Overlaps: Region overlaps between BTBR genetic variants (Indels or SNPs) and significant DARs. See methods for permutation testing details. All pvalues were FDR corrected for multilple comparisons.

**Table S7 Annotation_ChIPSeeker**
Annotation of peak sets for AllConsensusPeaks and each DAR peak set using ChIPSeeker.

**TableS8. DARpromoterGenes_DEGlistOverlaps**
FDR pvalue: FDR corrected p-values (q<0.05) for overlap between genes with promoter DARs and differentially expressed genes (DEGs) from the Fluidigm Array that were categorized as either non-tolerized or tolerized in either C57 or BTBR, or as hyper-responsive in BTBR under either LPSx1 or LPSx2 conditions. Statistics are from one tailed Fisher’s exact tests.
Odds Ratio: Odds ratio from Fisher’s exact test.
Raw Counts Overlapping Genes: The number of genes shared between each list.
Overlapping Gene Symbols: Gene symbols for all genes in each overlap comparison.
Overlapping Gene Symbols: Ensembl gene IDs for all genes in each overlap comparison.
